# EMT induces cell-cycle-dependent changes of Rho GTPases and downstream effectors

**DOI:** 10.1101/2023.02.02.526815

**Authors:** Kamran Hosseini, Annika Frenzel, Elisabeth Fischer-Friedrich

## Abstract

Epithelial-mesenchymal transition (EMT) is a key cellular transformation for many physiological and pathological processes ranging from cancer over wound healing to embryogenesis. Changes in cell migration, cell morphology and cellular contractility were identified as hallmarks of EMT. These cellular properties are known to be tightly regulated by the actin cytoskeleton. EMT-induced changes of actin-cytoskeletal regulation were demonstrated by previous reports of cell-cycle-dependent changes of actin cortex mechanics in conjunction with characteristic modifications of cortex-associated f-actin and myosin. However, at the current state, the changes of upstream actomyosin signalling that lead to corresponding mechanical and compositional changes of the cortex are not well understood. In this work, we show in breast epithelial cancer cells MCF-7 that EMT results in characteristic changes of the cortical signalling of Rho-GTPases Rac1, RhoA and RhoC and downstream actin regulators cofilin, mDia1 and Arp2/3. In the light of our findings, we propose that cell-cycle-dependent EMT-induced changes in cortical mechanics rely on two hitherto unappreciated signalling paths - i) a cell-cycle-dependent interaction between Rac1 and RhoC and ii) an inhibitory effect of Arp2/3 activity on cortical association of myosin II.

## 1. Introduction

Epithelial mesenchymal transition (EMT) is a cellular transformation of epithelial cells that entails the loss of apical-basal cell polarity and intercellular adhesion in combination with a gain of mesenchymal cell traits, see Fig. 1a [1–4]. EMT was linked to the initiation of metastasis and bad cancer prognosis through the acquisition of aggressive traits in cancer cells of epithelial origin [1–4]. In particular, EMT was reported to be connected to enhanced cell migration and cell proliferation in metastatic cancer cells [1–5].

**Figure 1.**
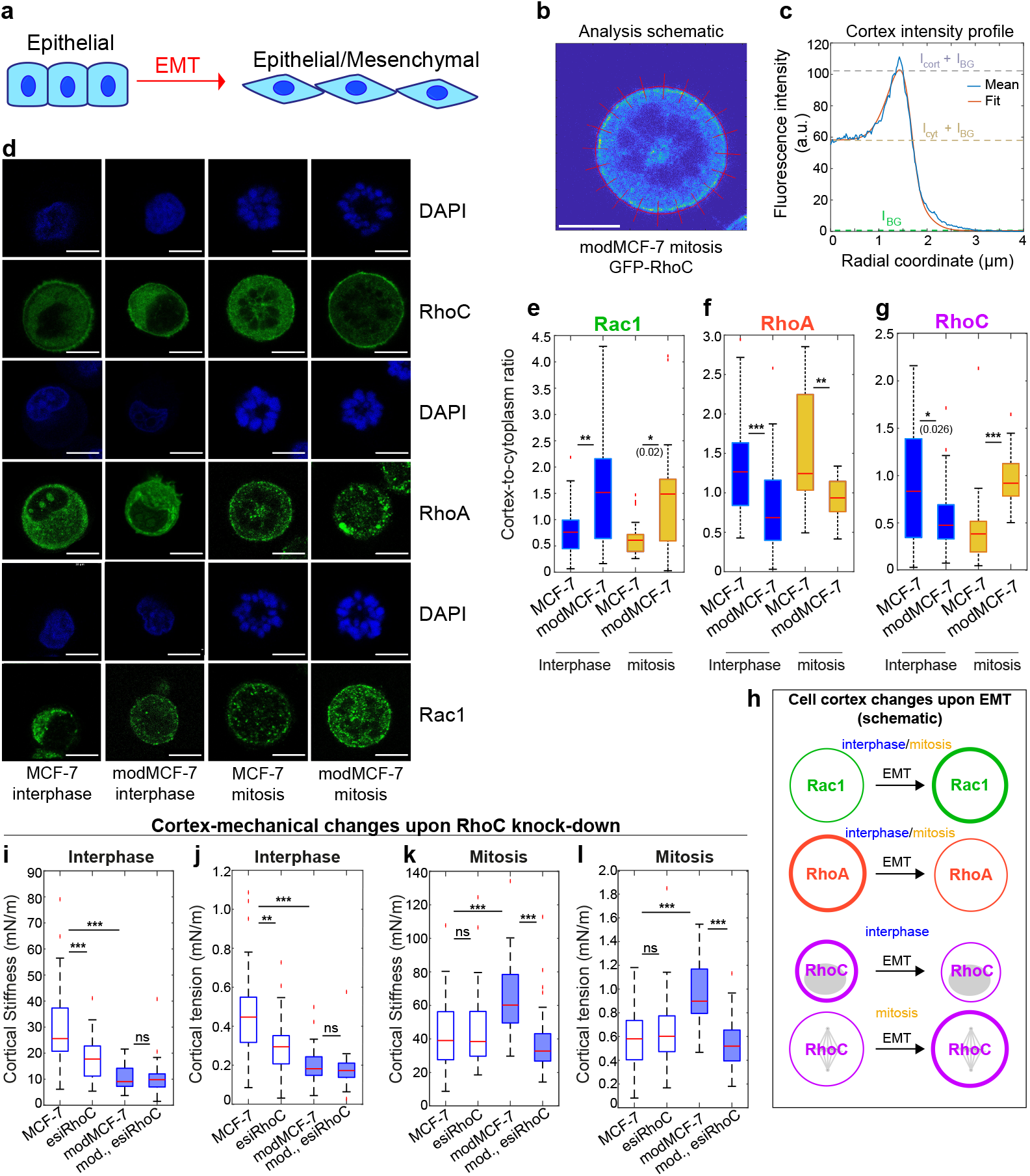
EMT induces characteristic changes of the cortical abundance of Rho GTPases RhoA, Rac1 and RhoC in MCF-7 cells. a) Schematic of EMT-induced changes of cell morphology and adhesion. b,c) Image analysis of confocal images of immunostained cell cross sections to extract the cortex-to-cytoplasm ratio. b) Exemplary picture of RhoC immunostaining fluorescence profile of the equatorial cross-section of an EMT-induced mitotic cell including elements of image analysis. Scale bar: 10 *μ*m. c) Mean radial fluorescence intensity profile of picture (blue curve) along radial lines shown in panel c. The fitted intensity profile, *Ism*(*r, p*) is shown in orange, see Materials and Methods. d) Representative confocal images of suspended interphase cells and STC-arrested mitotic cells in control and EMT-induced conditions. Cells were fixed and DAPI-stained for DNA (blue) and immunostained for Rac1/RhoA/RhoC (green), see Materials and Methods. Scale bar: 10 *μ*m. e-g) Cortex-to-cytoplasm ratio of RhoA, Rac1 and RhoC inferred from immunofluorescence staining as shown in panel b before and after EMT. h) Schematic of changes of cortical association of Rac1, RhoA and RhoC upon EMT. i-l) RhoC knockdown elicits cortical softening and tension reduction in the actin cortex in pre-EMT interphase MCF-7 cells (i-j, white boxplots) and post-EMT mitotic MCF-7 cells (k-l, blue-shaded boxplots). Post-EMT cells are referred to as modMCF-7. Number of cells analyzed: e: MCF-7 interphase n=34, modMCF-7 interphase n=32, MCF-7 mitosis n=31, modMCF-7 mitosis n=31. f: MCF-7 interphase n=36, modMCF-7 interphase n=28, MCF-7 mitosis n=31, modMCF-7 mitosis n=31. g: MCF-7 interphase n=44, modMCF-7 interphase n=43, MCF-7 mitosis n=30, modMCF-7 mitosis n=32. i-j: MCF-7 n=39, esiRhoC n=37, modMCF-7 n=37, esiRhoC n=39, k-l: MCF-7 n=27, esiRhoC n=29, modMCF-7 n=29, esiRhoC n=28. Measurements represent at least two independent experiments. n.s.: *p >* 0.05, *\** : *p <* 0.05, *\*\** : *p <* 0.01, *\* * \** : *p <* 0.001.

The actin cytoskeleton is a major regulator of cell mechanics, cell shape and cellular force generation. Thereby, the actin cytoskeleton constitutes a key player in cancer-related changes of cell migration and cell division [6–8]. Consistent with this, it was found that EMT causes major changes in the actin-cytoskeleton [5, 9, 10].

Rho GTPases are known to be essential regulators of the actin cytoskeleton. We and others showed that EMT is associated with characteristic changes in the activation of Rho GTPases as judged by the abundance of its active GTP-bound forms [5, 11–13]. In particular, we reported a decrease of total RhoA-GTP and an increase of total Rac1-GTP upon EMT in MCF-7 breast epithelial cells. Furthermore, the increased expression of the Rho GTPase RhoC was associated to enhanced metastasis in several cancer types [14]. In addition, RhoC signalling was featured to be essential for EMT [13–16].

Previously, we reported characteristic EMT-induced cell-cycle-dependent changes of actin cortex mechanics in rounded cells of diverse epithelial cancer cell lines originating from breast, lung, prostate and skin tissue indicating that this EMT-induced cell-mechanical change is a widely conserved feature in cells of diverse tissue origin [5, 17, 18]. Cortex-mechanical changes were entailing cortical softening and contractility reduction in interphase but a cortical stiffening and contractility increase upon EMT in mitosis. Concomitantly, we found EMT-induced changes of cortical actin and myosin II with reduced cortical myosin in interphase and increased cortical actin in mitosis [5].

While associated EMT-induced changes in Rho GTPases signalling provide a viable hypothesis for downstream changes of cortical actin and myosin, the details of EMT-induced changes in cortical signalling remain elusive. In particular, it is unclear how cortical mechanics is affected in an opposite way in interphase and mitosis, as none of the downstream actomyosin effectors of Rho GTPases is known to induce mechanical changes to the cortex that depend on the cell cycle stage [19, 20].

With this work, we aim to deepen our understanding of EMT-induced changes in cortical signalling, cortical composition and cortical mechanics with a focus on the differences between interphase and mitosis. To this end, we quantify EMT-induced changes of Rho GTPases RhoA, RhoC and Rac1 which were previously linked to EMT. Furthermore, we investigate as actin-regulating downstream targets formin, Arp2/3 and cofilin. In particular, we provide a quantitative analysis of EMT-induced changes of cortical protein localization in non-adherent cells in combination with changes of cortical mechanics and protein expression. In light of our results, we propose that two hitherto unappreciated signalling mechanisms at the cortex are at the heart of the cell-cycle dependent changes in cell mechanics - i) an interaction between Rac1 and RhoC and ii) an inhibitory effect of Arp2/3 activity on myosin II cortical localization.

## 2. Results

To investigate the effects of EMT on actin-cortical signalling and mechanics, we chose to work with the breast epithelial cancer cell line MCF-7 which exhibits epithelial cell traits in control conditions. We induced EMT in these cells via an established method (see e.g. [5, 21–25]) that entails a 48 hours treatment with the tumor promoter 12-O-tetradecanoylphorbol-13-acetate (TPA) at 100 nM, see Materials and Methods. We and others showed previously that in response to this treatment, MCF-7 cells display an EMT-characteristic protein expression change, corresponding cell-morphological changes towards a mesenchymal-like phenotype, as well as increased proliferation and migration, see e.g. [5, 24, 26]. We note that EMT-transformed cells will be referred to as modMCF-7 cells throughout this manuscript.

Previous research has shown that activation of Rho GTPases is connected to their association with the plasma membrane [27]. Furthermore, the activation of downstream effectors cofilin, formin and Arp2/3 was shown to be linked to f-actin binding [28–31]. Correspondingly, association of these proteins to cortical f-actin is a measure of their activity at the actin cortex. Therefore, one prevalent strategy of this study is to quantify changes of the relative amount of cortex-associated cortical regulators upon EMT as a readout of EMT-induced changes in cortical signalling. Following previous studies [5, 18, 19, 32, 33], we worked with rounded, non-adherent cells since this has the advantage that cell shapes are spherical in both epithelial and EMT-transformed conditions with a largely uniform actin cortex. In this way, a meaningful comparative analysis of cortical protein association between epithelial and the mesenchymal-like cells becomes possible.

For the measurement of cortical protein association, immunostaining of the cortical regulator under consideration was combined with fluorescent DNA staining (DAPI or Hoechst) which allowed to identify cells to be in an interphase or mitotic stage, see Materials and Methods. For the measurement of cortical regulators in mitotic cells, the fraction of mitotic cells was enriched through mitotic arrest induced by co-incubation with S-trityl-L-cysteine (STC), see Materials and Methods. Using a previously established image analysis scheme, we analysed confocal images of immunostaining fluorescence intensities to infer the cellular outline and the averaged cortical fluorescence profile along the radial coordinate, i.e. orthogonal to the cell boundary, see Fig. 1b and [5, 18]. The averaged radial fluorescence intensity was then used to derive the cortex-to-cytoplasm ratio of protein localization in the cells by calculating the ratio of the integrated cortical fluorescence normalized by the cytoplasmic fluorescence intensity, see Fig. 1c and Materials and Methods and [5, 18].

### 2.1. Rho GTPases change their cortical association upon EMT in a cell-cycle-dependent manner

To investigate whether cortical association of Rho GTPases changes through EMT, we quantified the cortex-to-cytoplasm ratio of RhoA, RhoC and Rac1 in cells with and without EMT induction. To this end, we performed confocal imaging of the equatorial cross section of suspended interphase or STC-arrested mitotic cells which were immunostained for either of the Rho GTPases under consideration, see Fig. 1d and Materials and Methods. Quantitative analysis shows that the cortex-to-cytoplasm ratio of Rac1 increases upon EMT both in interphase and mitosis (Fig. 1d,e). By contrast, the cortex-to-cytoplasm ratio of RhoA decreases through EMT (Fig. 1d,f). We conclude that cortical association of Rac1 and RhoA follows the EMT-induced quantitative change of GTP-bound Rac1 and RhoA in whole-cell-lysates of MCF-7 cells [5]. The cortex-to-cytoplasm ratio of RhoC changes in a cell-cycle-dependent manner upon EMT. While cortical RhoC goes down in interphase, we see an increase of cortical RhoC in mitosis (Fig. 1d,g). Our results on the effect of EMT on cortical signalling of Rho GTPases is summarized in Fig. 1h.

We further asked about the influence of Rho GTPases on cortical mechanics. For this purpose, we relied on cortex-mechanical measurements with an established cell confinement setup based on oscillatory cell-squishing with the cantilever of an atomic force microscope (AFM). We chose a deformation frequency of 1 Hz. This assay was previously shown to provide a readout of cortical stiffness, cortical tension as well as a characterization of the viscoelastic nature of the cortex quantified by the phase shift between stress and strain [5, 17, 18, 34]. (The phase shift takes values between 0 − 90° with lower values corresponding to a more solid-like response). In particular, we previously showed that Rac1 signalling was linked to a decrease in cortical stiffness and contractility in interphase cells but to an increase of cortical stiffness and contractility in mitotic cells with a stronger effect in post-EMT cells [5]. This is in agreement with our here reported finding of increased cortical Rac1 association post-EMT (Fig. 1e). On the other hand, we previously found that RhoA signalling increases cortical stiffness and contractility in particular in pre-EMT cells [5]. Again, the bigger mechanical effect pre-EMT is in agreement with our current observation of higher cortical RhoA association pre-EMT (Fig. 1f).

The effect of RhoC on cortical mechanics has to our knowledge not been reported previously. Using the AFM-based cell confinement assay, we measured cortical mechanics with and without RhoC knock-down via RNA interference in pre- and post-EMT conditions, see Fig. 1i-l, S1 and Materials and Methods. We find that similar to RhoA, RhoC signalling increases cortical contractility and stiffness, see Fig. 1i-l. However, this effect is restricted to pre-EMT interphase cells and post-EMT mitotic cells. It is plausible that the absence of an effect in these conditions is linked to low abundance of RhoC at the cortex (Fig. 1g). Furthermore, RhoC knock-down increases the phase shift in pre-EMT interphase conditions indicating that RhoC signalling contributes to the solid-like nature of the cortex in interphase, see Fig.

### 2.2. Rac1 and RhoC mutually affect their cortical association in a cell-cycle-dependent manner

The previously reported finding that Rac1 activity affects cortical mechanics opposite in interphase and mitosis provides a clue that the signalling of Rac1 might be at the heart of the cell-cycle dependence of cytoskeletal changes upon EMT. However, currently it is unclear how cortical Rac1 signalling can act in a manner that is qualitatively different in interphase and mitosis. In particular, it surprised us that Rac1 would make a strong contribution to cortical contractility in mitosis in post-EMT conditions given that Arp2/3 activity increase downstream of Rac1 is expected to diminish cortical contractility, see Fig. 6 and [20, 35]. Furthermore, previous reports showed that RhoA is at the heart of cortical contractility in mitosis [36]. While RhoA activity and cortical association is low in post-EMT MCF-7 cells (Fig. 1f and [5]), we note that RhoC signalling is similar to RhoA. Therefore, RhoC signalling might step in for RhoA signalling after EMT during mitosis.

Following this line of thought, we asked whether Rac1 might increase cortical contractility in post-EMT mitosis via (direct or indirect) activation of RhoC. To test this hypothesis, we monitored changes in cortical association of RhoC upon knock-down of Rac1 in pre- and post-EMT conditions judged by fluorescence intensity of RhoC immunostaining (Fig. 2a). Obtained confocal images of equatorial cross-sections were used for image analysis in all conditions. We find that inferred cortex-to-cytoplasm ratios of RhoC increase upon knock-down of Rac1 in rounded interphase cells with a stronger effect in post-EMT conditions (Fig. 2a,c). By contrast, the cortex-to-cytoplasm ratio of RhoC decreases upon knock-down of Rac1 in mitosis with a stronger effect in post-EMT conditions (Fig. 2a,d). We conclude that Rac1 signalling increases cortical association of RhoC in mitosis but diminishes cortical association in interphase in MCF-7 cells (Fig. 2g).

**Figure 2.**
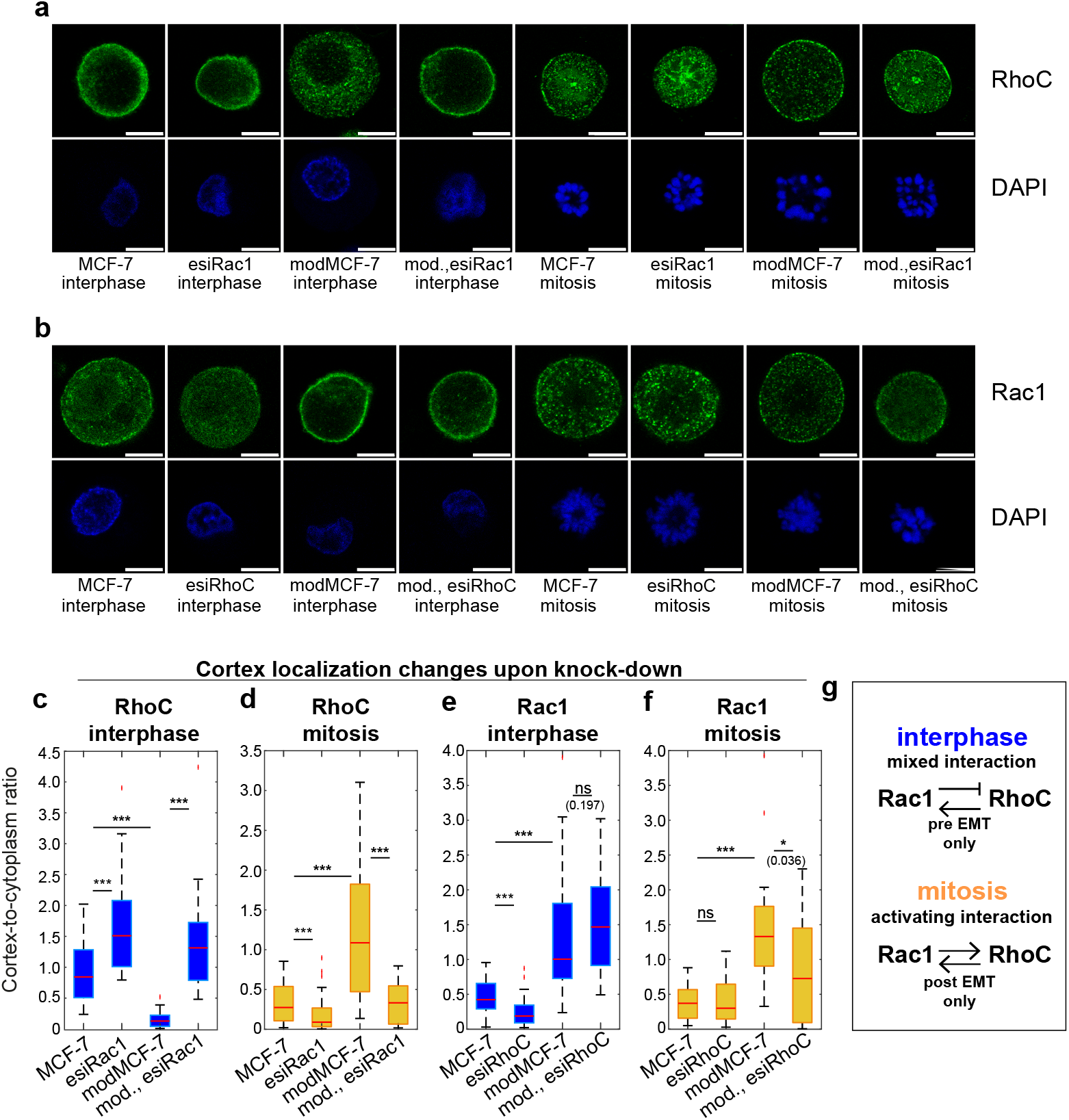
Rho-GTPases Rac1 and RhoC mutually affect their abundance at the cortex in a cell-cycle-dependent manner. a) Representative confocal images of RhoC (immunostained, green) and DNA (DAPI, blue) with and without Rac1 knock-down in suspended interphase cells and STC-arrested mitotic cells in pre-EMT (MCF-7) and post-EMT (modMCF-7) conditions. Scale bar: 10 *μ*m. b) Representative confocal images of Rac1 (immunostained, green) and DNA (DAPI, blue) with and without RhoC knock-down in suspended interphase cells and STC-arrested mitotic cells in pre-EMT (MCF-7) and post-EMT (modMCF-7) conditions. Scale bar: 10 *μ*m. c-f) Changes of cortical association of RhoC and Rac1 upon knock-down of the other protein, i.e. Rac1 or RhoC, respectively. Cortical association of either protein was quantified by its cortex-to-cytoplasm ratios which was inferred from immunofluorescence staining as shown in panel a and b before and after EMT. g) Schematic summary of Rac1 and RhoC mutual interactions in interphase and mitosis. Post-EMT cells are referred to as modMCF-7. Number of cells analyzed: c: MCF-7 n=20, esiRac1 n=20, modMCF-7 n=20, esiRac1 n=20, d: MCF-7 n=19, esiRac1 n=22, modMCF-7 n=19, esiRac1 n=19, e: MCF-7 n=25, esiRhoC n=21, modMCF-7 n=20, esiRhoC n=24, f: MCF-7 n=24, esiRhoC n=26, modMCF-7 n=22, esiRhoC n=24. Measurements represent at least two independent experiments. n.s.: *p >* 0.05, *\** : *p <* 0.05, *\*\** : *p <* 0.01, *\* * \** : *p <* 0.001.

To test whether in return also RhoC signalling influences Rac1, we performed immunostaining of Rac1 with and without RhoC knock-down in pre- and post-EMT conditions in interphase and mitosis, see Fig. 2b. We find that inferred cortex-to-cytoplasm ratios of Rac1 decrease upon knock-down of RhoC in pre-EMT interphase cells and in post-EMT mitotic cells (Fig. 2e,f). In all other conditions, there is no significant effect on RhoC cortical association (Fig. 2e,f). We conjecture that the signalling from RhoC to Rac1 is restricted to pre-EMT interphase and post-EMT mitosis due to the low cortical representation of RhoC in post-EMT interphase and pre-EMT mitotic conditions, see Fig. 1g,h. With this explanation approach, our data are consistent with an in general activating effect of RhoC on Rac1 (Fig. 2g).

In previous work, the active forms of RhoA and Rac1 were shown to affect each other through mutually inhibitory interactions in breast epithelial cells [37]. This is consistent with the EMT-induced switch-like change from a state of high RhoA and low Rac1 activation to a state of low RhoA and high Rac1 activation [5]. The cell-cycle dependent interaction between Rac1 and RhoC has been to our best knowledge unknown so far.

### 2.3. Cortical cofilin association increases through EMT

We went on to ask how EMT-induced changes in Rho GTPases signalling affect downstream cortical regulators. We first investigated EMT-induced changes of cofilin cortical association. Cofilin is known to promote the depolymerization of the actin cortex through severing of actin fibers [38]. In the context of cancer, cofilin activity at the cortex has been suggested to be a main factor in f-actin turnover thus playing a key role in cancer cell migration and invasion [39]. Cofilin becomes deactivated through phosphorylation mediated by Lim kinases [39] and phosphorylated cofilin was shown to not interact with f-actin [28]. Correspondingly, cortex-bound cofilin can be interpreted as active cofilin. Quantifying amounts of total cofilin (CFL1) and phospho-cofilin (phospho-CFL1 (Ser3)) via western blotting from lysates of adherent cells, we find a trend of a shallow increase of total cofilin (only interphase) and a decrease of phospho-cofilin upon EMT, see Fig. 3a-c and Materials and Methods. Taken together, this points at an increase of the active non-phosphorylated form of cofilin upon EMT in MCF-7.

**Figure 3.**
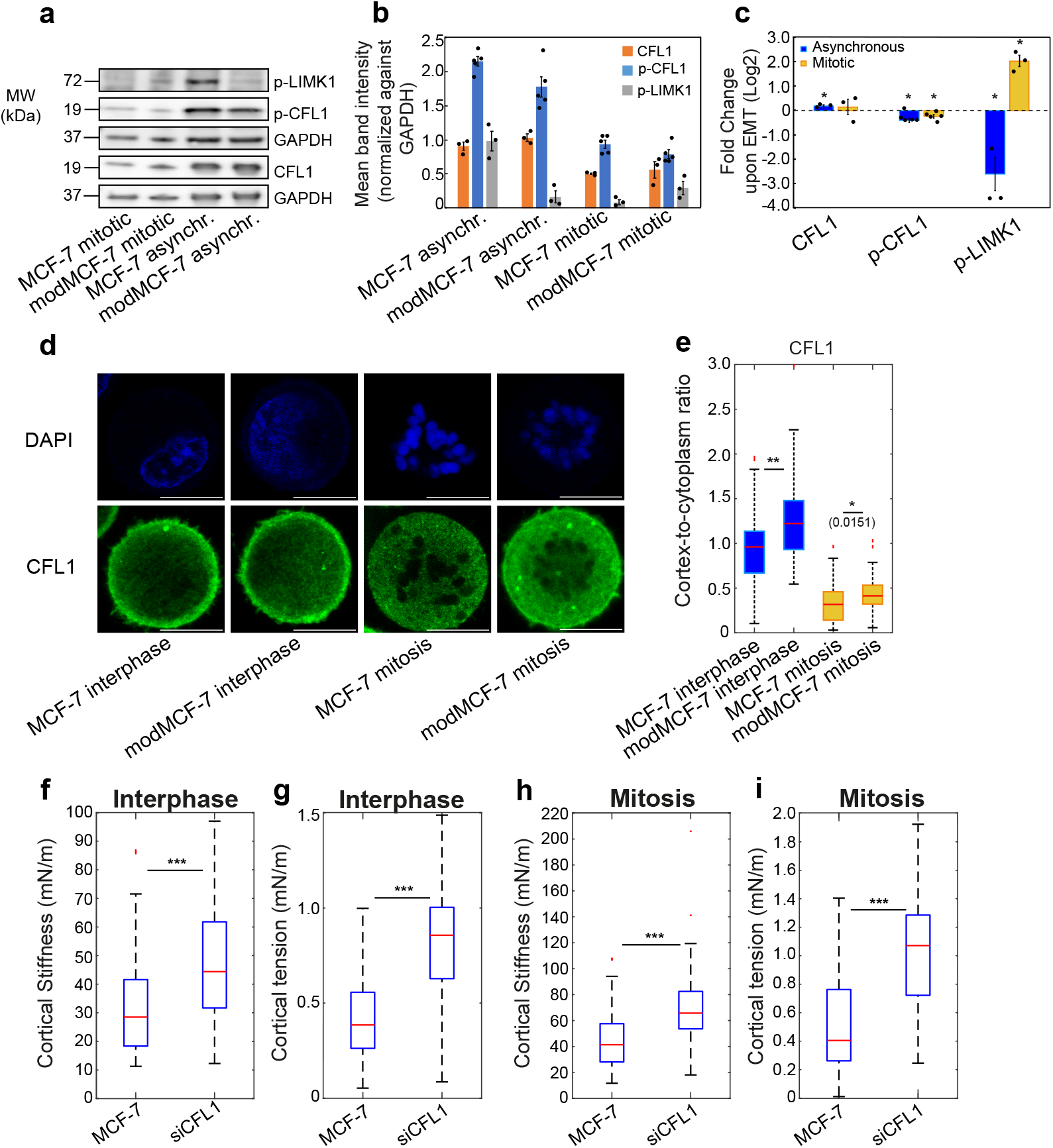
Actin cortical regulators LIMK1 and cofilin (CFL1) influence actin cortical mechanics and change their cortical representation upon EMT. a) Exemplary western blots for p-LIMK1, p-CFL1 and CFL1 expression in MCF-7 cells in control and EMT-induced conditions. b) Bar charts of normalized quantities of the p-LIMK1 (active form), p-Cofilin (inactive form) and total cofilin western blots. Normalization was done against GAPDH bands. Error bars represent standard error of the mean. c) Fold changes of p-LIMK1, p-CFL1 and total CFL1 upon EMT from western blots. Individual data points are depicted in black. Error bars represent standard error of the mean. d) Representative confocal images of suspended interphase cells and STC-arrested mitotic cells upon EMT, fixed and stained for DAPI (blue) and cofilin (green). Scale bar: 10 *μ*m. e) Cortex-to-cytoplasm ratio of cofilin inferred from immunofluorescence staining as shown in panel d before and after EMT. f-i) CFL1 knockdown elicits cortical stiffening and tension rise in the actin cortex in interphase (f-g) and mitotic (h-i) MCF-7 cells. Post-EMT cells are referred to as modMCF-7. Number of cells analyzed: e: MCF-7 interphase n=38, modMCF-7 interphase n=38, MCF-7 mitosis n=46, modMCF-7 mitosis n=44. f-g: MCF-7 n=49, siCFL1 n=49, h-i: MCF-7 n=46, siCFL1 n=50. Measurements represent at least two independent experiments. n.s.: *p >* 0.05, *\** : *p <* 0.05, *\*\** : *p <* 0.01, *\* * \** : *p <* 0.001.

To assess cortical association of cofilin, we performed also cofilin-immunostaining of rounded cells, see Fig. 3d. We find that the cortex-to-cytoplasm ratio of cofilin is elevated upon EMT indicating an EMT-mediated increase of cortical cofilin activity, see Fig. 3d,e. This finding is in agreement with an increase of active cofilin as suggested by western blotting.

Immunostaining of the cofilin upstream regulator phospho-Limk1 (phospho-LIMK1 (Thr508)) shows no cortical association but cytoplasmic localization in agreement with previous findings, see Fig. S3a and [40]. Quantifying phospho-Limk1 abundance in whole-cell lysates via western blotting, we find an EMT-induced decrease in interphase (asynchronous cell population) in accordance with the observed concomitant decrease of phospho-cofilin, see Fig. 3a-c and Materials and Methods. In mitosis, phospho-Limk1 amounts are very low and show a trend of decrease upon EMT which is, surprisingly, in disagreement with the EMT-induced trend of phospho-cofilin (Fig. 3a-c). We speculate that this apparent inconsistency may be attributed to the previously reported modified activation scheme of Limk1 in mitosis, where hyperphosphorylation rather than phosphorylation at Thr508 is at the heart of Limk1 activation [41]. This observation features phospho-Limk1 (Thr508) as an unsuitable readout of cofilin phosphorylation activity in mitosis.

Investigating the effect of cofilin on cortical mechanics, we find that cofilin knock-down through RNA interference changes cortical mechanics, see Fig. 3f-i and Fig. S3e-h. Both cortical tension and stiffness increase in interphase and mitosis, see Fig. 3f-i. In addition, the phase shift and therefore the fluid-like nature increased mildly upon knockdown in interphase cells, see Fig. S3f. We conclude that increased cortical association of cofilin in EMT-induced cells contributes to a trend of decreased cortical contractility and stiffness.

We note that Chugh et al. [19] previously reported a tension increase upon CFL1-knockdown in interphase HeLa cells in agreement with our findings. However, the authors reported by contrast a tension decrease upon CFL1 knock-down in mitosis opposite to our findings in MCF-7. This apparent discrepancy might be rooted in the non-monotonous dependence of cortical tension on actin filament length as was proposed by the same study [19]. According to this idea, increased actin filament length through cofilin knock-down can either increase or decrease cortical tension depending on the initial state of the cortex. Large differences in cortical tension values between mitotic MCF-7 cells and mitotic HeLa cells make different cortical configurations in mitosis for the two cell lines additionally plausible.

### 2.4. The actin nucleator mDia1 shows cell-cycle-dependent changes of cortical association upon EMT

In order to further understand EMT-induced changes of cortical composition and mechanics [5], we addressed how actin nucleators are affected upon EMT downstream of Rho GTPases. Cortical actin is polymerized by formins and Arp2/3. We will first focus on the influence of the former. Previous studies have shown that formin activity has a major influence on cortical mechanics [19, 20, 42, 43]. We confirmed this finding in rounded MCF-7 cells with our AFM-based cell confinement setup showing that formin inhibition via 40 *μ*M SMIFH2 reduced cortical stiffness and contractility in MCF-7 cells in interphase and mitosis, see Fig. S4.

To further investigate how formin-mediated polymerization changes at the cortex upon EMT, we decided to focus on the formin representative mDia1 (also Diaph1) which together with the actin nucleator Arp2/3 polymerizes the majority of f-actin in the actin cortex [44]. mDia1 is activated downstream of RhoA, RhoB or RhoC through binding to the active form of the respective Rho GTPase [30]. The active form of mDia1 associates with f-actin [30, 31] and thus with the actin cortex.

Performing quantification of protein amounts in whole-cell lysates via western blots, we find that there is no significant change of expression of mDia1 upon EMT (Fig. 4a-c). We then went on to monitor cortical association of mDia1 via immunostaining and quantification of the cortex-to-cytoplasm ratio, see Fig. 4d,e. We find clear EMT-induced changes; in interphase cells, the cortex-to-cytoplasm ratio of mDia1 is reduced, see Fig. 4e, blue boxes. We conclude that cortical association of mDia1 is decreased upon EMT in agreement with our finding of reduced presence of RhoA and RhoC at the cortex. In mitotic cells, on the other hand, the cortex-to-cytoplasm ratio of mDia1 is increased, see Fig. 4e, yellow boxes. This finding points at an increase of cortical mDia1 activity upon EMT in mitosis. We suggest that this effect is due to an EMT-induced rise of cortical activity of RhoC in mitosis overcompensating the effect of reduced RhoA activity in post-EMT mitotic cells, see Fig. 1h.

**Figure 4.**
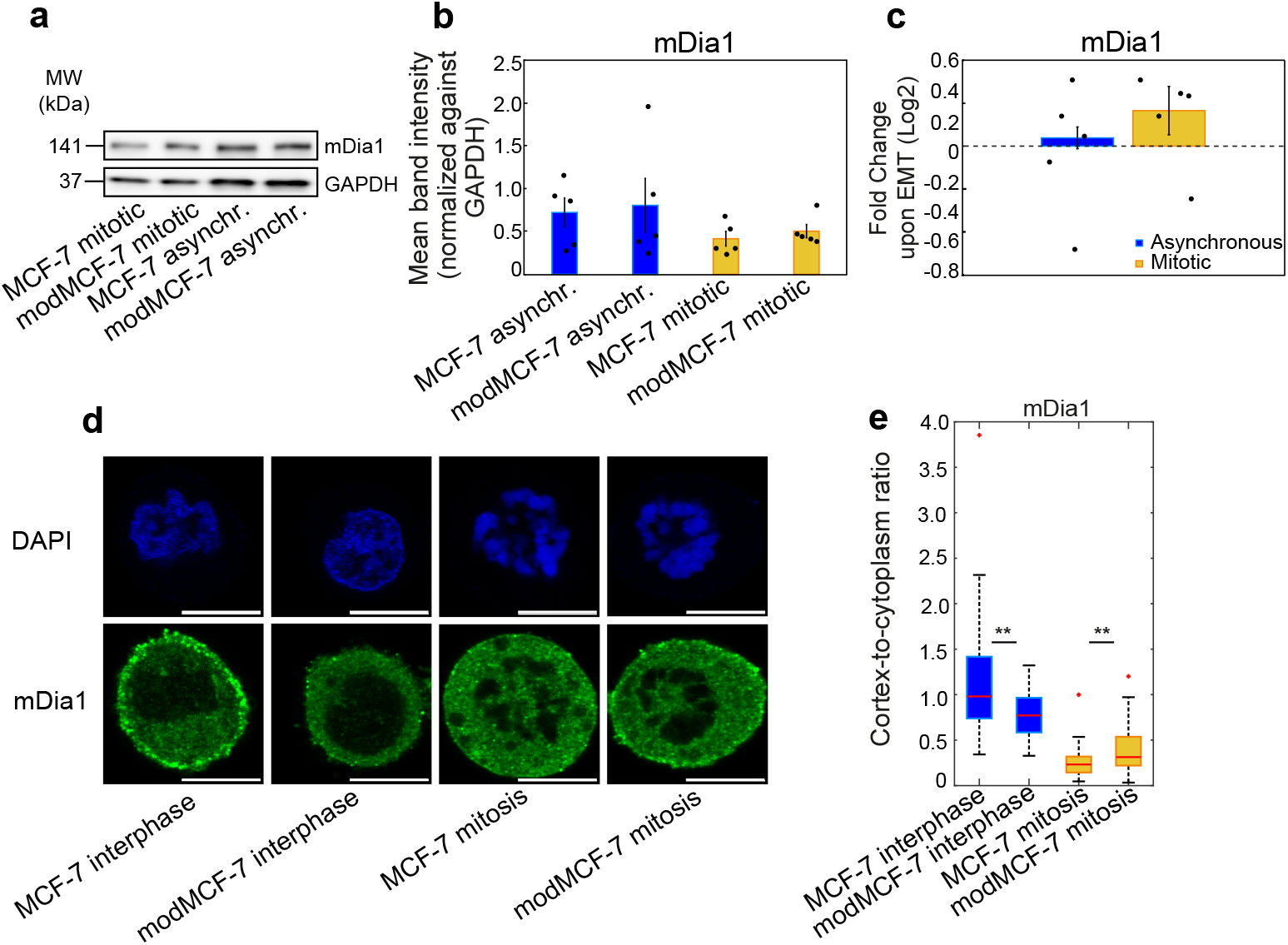
The actin nucleator and RhoA/C downstream effector mDia1 changes its cortical association upon EMT in a cell-cycle dependent manner. a) Exemplary western blots for mDia1 expression in MCF-7 cells in control and EMT-induced conditions. b) Bar charts of normalized quantities of mDia1 western blots. Normalization was done against GAPDH bands. Error bars represent standard error of the mean. c) Fold changes of mDia1 upon EMT induction from western blots. Individual data points are depicted in black. Error bars represent standard error of the mean. d) Representative confocal images of suspended interphase cells and STC-arrested mitotic cells upon EMT, fixed and stained for DAPI (blue) and mDia1 (green). Scale bar: 10 *μ*m. e) Cortex-to-cytoplasm ratio of mDia1 inferred from immunofluorescence staining as shown in panel d before and after EMT. Post-EMT cells are referred to as modMCF-7. Number of cells analyzed: e: MCF-7 interphase n=72, modMCF-7 interphase n=32, MCF-7 mitosis n=33, modMCF-7 mitosis n=36. Measurements represent at least two independent experiments. n.s.: *p >* 0.05, *\** : *p <* 0.05, *\*\** : *p <* 0.01, *\* * \** : *p <* 0.001.

Taken together, our results suggest that the observed EMT-induced cell-cycle-dependent changes of cortical mDia1 likely make an essential contribution to cell-cycle-dependent EMT-induced changes of cortex-associated actin and cortical mechanics.

### 2.5. The actin nucleator Arp2/3 increases its cortical association upon EMT

To further increase our understanding of changes in cortex-associated actin upon EMT as reported in [5], we also investigated EMT-induced changes of the second major actin nucleator beyond mDia1, namely the Arp2/3 complex [6, 44]. The Arp2/3 complex becomes activated by the Wasp family of proteins [29, 45]. Activated Wasp proteins promote binding of the Arp2/3 complex to f-actin [29, 46, 47] and thus its association to the actin cortex. Wasp proteins (WAVE) are activated downstream of Rac1 [48]. We therefore expect that our finding of EMT-induced increase of cortical Rac1 association (Fig. 1h) should give rise to a downstream increase cortical Arp2/3 association. Performing protein quantification in whole-cell lysates, we find that there are no significant expression changes of Arp2 upon EMT, see Fig. 5a-c. This indicates that there are no significant changes of the amount of the Arp2/3 complexes in MCF-7 cells upon EMT induction. To assess cortical association of the Arp2/3 complex, we performed immunostaining of Arp2 in rounded cells in interphase and mitosis, see Fig. 5d. Interestingly, in spite of a direct interaction between Arp2/3 and f-actin, a cortical enrichment of Arp2 is only visible in interphase cells, see Fig. 5d, lower row. Quantification of corresponding cortical association in interphase via the cortex-to-cytoplasm ratio indicates that Arp2/3 signalling at the cortex is enhanced through EMT, see Fig. 5e.

**Figure 5.**
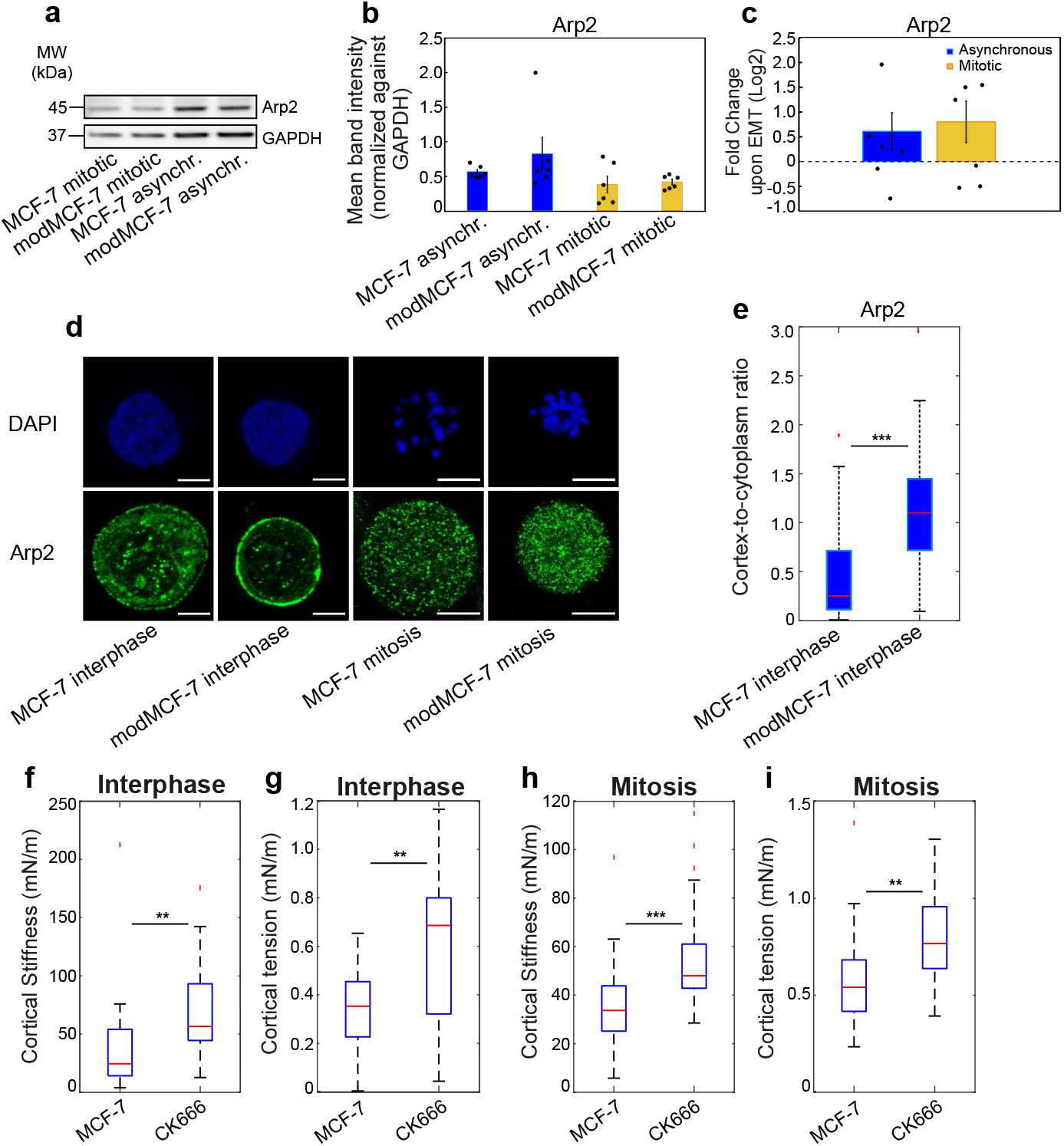
The actin nucleator and Rac1 downstream effector Arp2/3 is elevated at the cortex upon EMT in interphase. a) Exemplary western blots for Arp2 expression in MCF-7 cells in control and EMT-induced conditions. b) Bar charts of normalized quantities of Arp2 western blots. Normalization was done against GAPDH bands. Error bars represent standard error of the mean. c) Fold changes of Arp2 upon EMT from western blots. Individual data points are depicted in black. Error bars represent standard error of the mean. d) Representative confocal images of suspended interphase cells and STC-arrested mitotic cells upon EMT, fixed and stained for DAPI (blue) and Arp2 (green). Scale bar: 5 *μ*m. e) Cortex-to-cytoplasm ratio of Arp2 inferred from immunofluorescence staining as shown in panel d before and after EMT. Mitotic cells did not show a clear cortical Arp2 association and were therefore not quantified. f-i) Arp2 inhibition using 50 *μ*M CK666 elicits cortical softening and tension increase in the actin cortex in interphase (f-g) and mitotic MCF-7 cells (h-i). Post-EMT cells are referred to as modMCF-7. Number of cells analyzed: e: MCF-7 interphase n=47, modMCF-7 interphase n=48. f-g: MCF-7 n=24, CK666 n=24. h-i: MCF-7 n=27, CK666 n=24. Measurements represent at least two independent experiments. n.s.: *p >* 0.05, *\** : *p <* 0.05, *\*\** : *p <* 0.01, *\* * \** : *p <* 0.001.

Previous studies reported that Arp2/3 signalling reduces cortical tension in interphase and mitosis [32, 35]. This is counter-intuitive as Arp2/3 mediates actin polymerization and, thereby, could be expected to increase the amount of cortical myosin II substrate. To test the effect of Arp2/3 signalling on cortical tension in MCF-7 cells, we measured actin cortical mechanics with and without the Arp2/3 inhibitor CK666, see Fig. 5f-i and Fig. S5. We find that in agreement with previous results, cortical tension and stiffness increases upon Arp2/3 inhibition in interphase and mitosis, see Fig. 5f-i. By contrast, the phase shift did not change significantly upon Arp2/3 inhibition, see Fig. S5b,d. We conclude that our data suggest that increased Arp2/3 activity upon EMT downstream of increased Rac1 activity tends to reduce cortex stiffness and contractility independent of the cell cycle state.

## 3. Arp2/3 activity enhances cortical actin but reduces cortical association of myosin II

To deepen our understanding of cortical response to EMT-induced changes of cortex-associated Arp2/3, we investigated how cortex-associated actin and myosin change in response to Arp2/3 inhibition. For this purpose, we transfected MCF-7 cells with constructs expressing fluorescently labeled myosin regulatory light chain (MLC2) or fluorescently labeled actin (ACTB) as fluorescent reporters of cellular myosin II and actin localization, see Fig. 6a and Materials and Methods. Cell transfection was performed for suspended cells in interphase and mitosis as well as in pre- and post-EMT conditions, see Materials and Methods. Quantifying cortical association of myosin II and actin via confocal imaging of transfected live cells, we determined the cortex-to-cytoplasm ratio of actin and myosin II in conditions with and without Arp2/3 inhibition via the inhibitor CK666. As expected, we find that cortical actin association goes down upon Arp2/3 inhibition in all conditions, see Fig. 6b. By contrast, we observe that Arp2/3 inhibition increases myosin II association to the cortex in all conditions in spite of reduction of cortical actin, see Fig. 6c.

**Figure 6.**
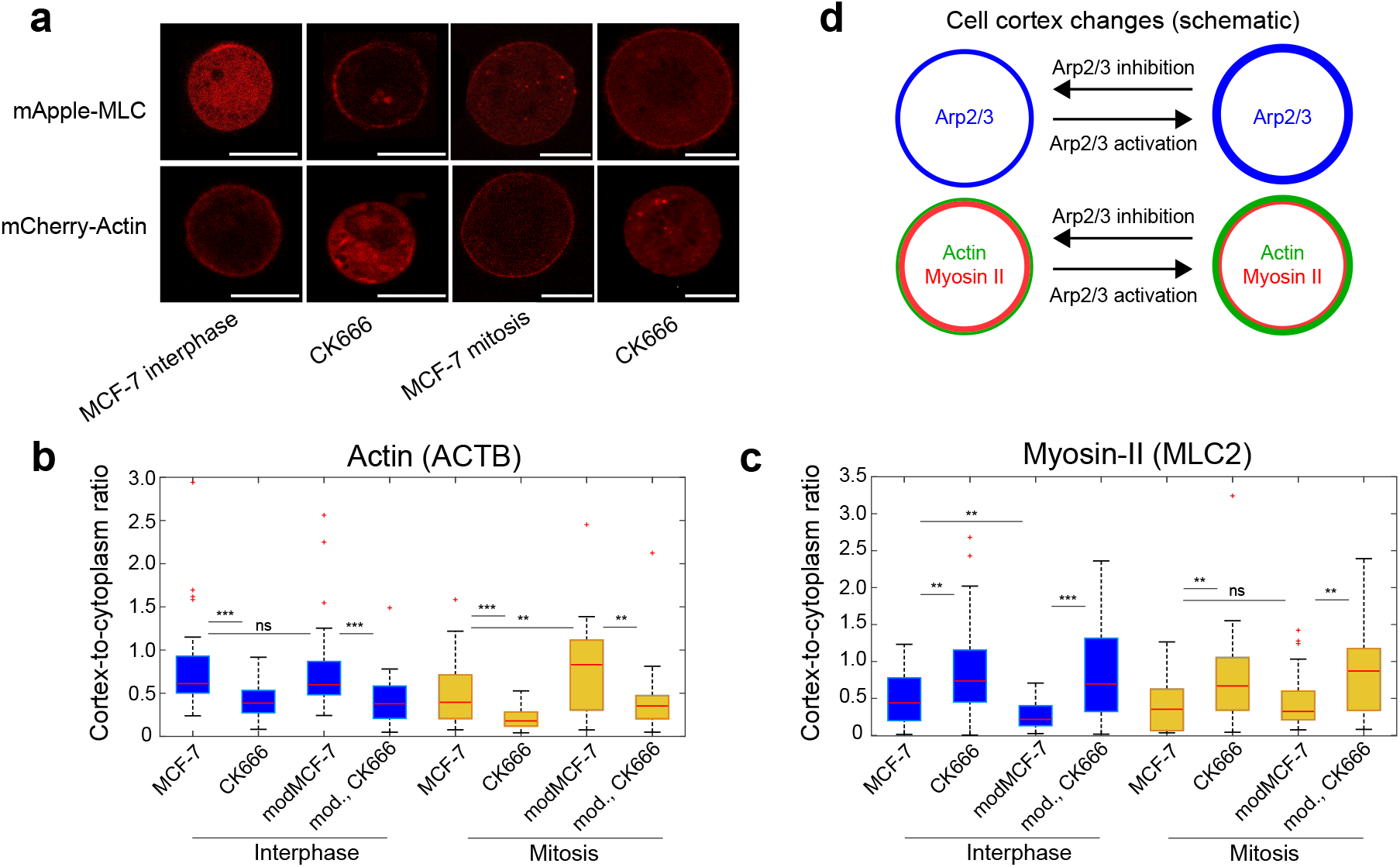
Activity of the Rac1 downstream effector Arp2/3 modulates actin and myosin association to the cortex. a) Representative confocal images of live suspended interphase cells and STC-arrested mitotic MCF-7 cells upon Arp2/3 inhibition with CK666, expressing mCherry-ACTB (bottom row) or mApple-MLC (top row). Scale bar: 10 *μ*m. b-c) Cortex-to-cytoplasm ratio of actin (b) and myosin (c) inferred from live images as shown in panel a before and after Arp2/3 inhibition with 50 *μ*M CK666. d) Schematic of cortical changes upon Arp2/3 activation/inhibition. Post-EMT cells are referred to as modMCF-7. Number of cells analyzed: b: interphase: MCF-7 n=35, CK666 n=34, modMCF-7 n=35, CK666 n=31, mitosis: MCF-7 n=31, CK666 n=30, modMCF-7 n=35, CK666 n=31, c: interphase: MCF-7 n=38, CK666 n=33, modMCF-7 n=38, CK666 n=35, mitosis: MCF-7 n=28, CK666 n=35, modMCF-7 n=27, CK666 n=30. n.s.: *p >* 0.05, *\** : *p <* 0.05, *\*\** : *p <* 0.01, *\* * \** : *p <* 0.001.

To further corroborate these findings, we double-checked the observed effect of Arp2/3 signalling in fixed cells using Phalloidin as a fluorescent reporter of f-actin and immunostaining of MYH9 as a fluorescent readout of myosin II localization, see Fig. S6. Again, we see that Arp2/3 inhibition leads to diminished cortical f-actin in combination with an increase in cortical myosin II confirming our results obtained from transfected live cells, see Fig. S6b,c.

We conclude that cortical Arp2/3 in addition to expected allocation and polymerization of cortical actin leads to diminished cortical association of myosin II. Increased cortex-associated Arp2/3 downstream of EMT-induced enhanced Rac1 signalling may therefore, at least in part, account for the observation of emergent reduced cortical tension and stiffness as well as reduced myosin II cortex-to-cytoplasm ratios upon EMT in interphase cells [5]. We note that a negative effect of Arp2/3 on myosin activity was previously reported within mouse oocytes [49].

## 4. Discussion

In this study, we investigated how actin cortex regulation is changed upon EMT in MCF-7 breast epithelial cancer cells. As previous work suggested that activity changes of the Rho GTPases are a game changer of cytoskeletal regulation upon EMT [5, 13, 37, 50–52], we focused on modulations of cortical signalling of RhoA, RhoC and Rac1 as well as on selected downstream effectors.

Signalling of cortical regulator proteins was assessed through immunostaining and subsequent confocal imaging of fixed MCF-7 breast epithelial cells in a rounded, non-adherent state with a largely uniform cortex. The magnitude of the cortex-to-cytoplasm ratio of the regulator protein under consideration was used as a quantitative readout of cortical signalling strength. In particular, we compared cortex-to-cytoplasm ratios in control and EMT-induced conditions in interphase and mitosis. Furthermore, for cortical regulators RhoC, formin, Arp2/3 and and cofilin, we identified their influence on cortical mechanics which was quantified by an established AFM-based cell confinement setup [5, 17, 18, 34].

In summary, we found that EMT reduces cortical association of RhoA but enhances cortical association of Rac1, while cortical RhoC is subject to a cell-cycle-dependent EMT-induced change, see Fig. 1h and Fig. 7. Interestingly, we discovered a hitherto unappreciated interaction between RhoC and Rac1 that likely contributes to RhoC activation in mitotic EMT-induced cells, see Fig. 2g. This interaction entails in particular a reduction of cortical RhoC in interphase but an increase of cortical RhoC in mitosis through Rac1 signalling.

**Figure 7.**
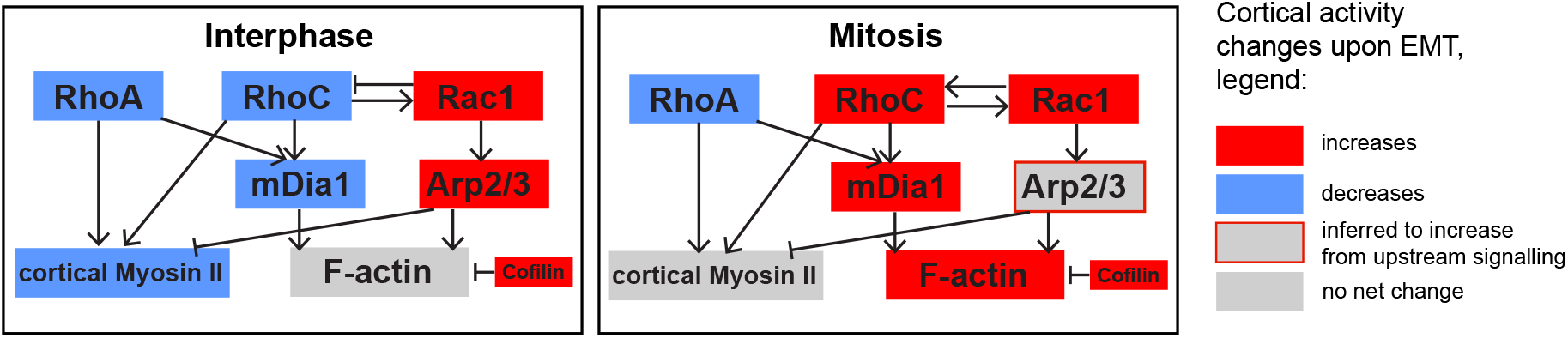
Schematic of cell-cycle-dependent EMT-induced changes of cytoskeletal signalling in MCF-7 breast epithelial cancer cells as suggested by our data. Red boxes indicate proteins whose cortical signalling increases through EMT. Blue boxes indicate proteins whose cortical signalling decreases through EMT. Grey boxes indicate proteins that show no net change in their cortical signalling upon EMT. In the signalling network, pointed black arrows represent activating signalling while flat black arrows indicate inhibiting signalling.

Downstream of Rho GTPases, we found that also the cortical signalling of the formin mDia1 is affected in a cell-cycle-dependent manner by EMT. The corresponding decrease of mDia1 at the EMT-transformed interphase cortex can be attributed to decreased RhoA and RhoC signalling. On the other hand, EMT-related increase of mitotic RhoC signalling can account for increased mDia1 at the mitotic post-EMT cortex, see Fig. 7. Furthermore, we find an EMT-induced increase in Arp2/3 and cofilin signalling at the cortex in both interphase and mitosis, see Fig. 7.

Taken together, our study indicates that actin nucleation at the interphase cortex is promoted upon EMT through upregulation of Arp2/3, but diminished through downregulation of mDia1 and enhanced cofilin signalling at the cortex. All in all, these signalling changes may give rise to the observed absence of a net change of cortex-associated actin upon EMT in interphase, see Fig. 6b and [5]. Furthermore, myosin activity at the interphase cortex is diminished through reduced RhoA and RhoC signalling (via Rock) and via increased Rac1 signalling (mediated downstream by Arp2/3, see Fig. 6d). The combined effects can account for the observed net decrease of cortex-associated myosin upon EMT and can be causative to cortical stiffness and contractility reduction, see Fig. 7.

In mitotic cells, EMT increases cortical actin nucleation through enhanced cortical RhoC signalling (e.g. via mDia1) and Rac1 signalling (via Arp2/3). On the other hand, EMT decreases cortical actin through reduced cortical RhoA signalling and enhanced cofilin signalling. The integration of all signalling changes can account for the observed net increase of cortical actin upon EMT in mitosis, see Fig. 6d and [5].

Further, EMT promotes myosin activity at the mitotic cortex through enhanced RhoC signalling, but diminishes it through reduced RhoA signalling and via increased Rac1 signalling (mediated downstream by Arp2/3 signalling, see Fig. 6d). Through the combination of these opposite effects, the net change of cortical myosin II may vanish as was observed in MCF-7 cells, see Fig. 7.

In conclusion, we find that EMT induces complex modifications in actin-cytoskeletal signalling through a combination of changes in the signalling of Rho GTPases and downstream effectors such as cofilin, Arp2/3 and mDia1. The integration of all partly opposing effects give rise to an emergent change of actin and myosin at the cortex. In particular, our findings shed further light on how cell-cycle-dependent differences emerge in cortical composition and mechanics. Finally, we note that our study provides a cellular EMT fingerprint of rounded cells that may be relevant for cancer diagnostic approaches in particular for those that rely on isolated cells such as FACS-related assays or deformability flow cytometry approaches [53–55].

## 5. Materials and Methods

### 5.1. Cell culture

The cultured cells were maintained as follows: MCF-7 cells were grown in RPMI-1640 medium (PN:2187-034, life technologies) supplemented with 10% v/v fetal bovine serum, 100 *μ*g/mL penicillin, 100 *μ*g/mL streptomycin (all Invitrogen) at 37°C with 5% CO2. In MCF-7 cells, EMT was induced by incubating cells in medium supplemented with 100 nM 12-O-tetradecanoylphorbol-13-acetate (TPA) (PN:P8139, Sigma) for 48 h prior to measurement [26]. Arp2/3 inhibition was performed by 30 minutes treatment with 50 *μ*M CK666 (PN:SML0006, Sigma). mDia1 inhibition was performed by 30 minutes treatment with 40 *μ*M SMIFH2, which inhibits all formins.

### 5.2. AFM measurement of cells

#### Experimental setup

To prepare mitotic cells for AFM measurements, approximately 10,000 cells were seeded in a cuboidal silicon cultivation chamber (0.56 *cm*^2^ area, from cutting ibidi 12-well chamber; ibidi, Gräfelfing, Germany) that was placed in a 35 mm cell culture dish (fluorodish FD35-100, glass bottom; World Precision Instruments, Sarasota, FL) 1 day before the measurement so that a confluency of ∼30% was reached on the measurement day. Mitotic arrest was induced by supplementing S-trityl-L-cysteine (Sigma-Aldrich) 2-8 hours before the measurement at a concentration of 2 *μ*M. For measurement, mitotic-arrested cells were identified by their shape. Their uncompressed diameter ranged typically from 18 to 23 *μ*m.

To prepare AFM measurements of suspended interphase cells, cell culture dishes (fluorodish FD35-100) and wedged cantilevers were plasma-cleaned for 2 minutes and then coated by incubating the dish at 37°C with 0.05 mg/mL poly(L-lysine)-polyethylene glycol dissolved in phosphate-buffered saline (PBS) overnight at 37°C (poly(L-lysine)(20)-g[3.5]-polyethylene glycol(2); SuSoS, Dubendorf, Switzerland) to prevent cell adhesion. Before measurements, cultured cells were detached by the addition of 0.05 trypsin-EDTA (Invitrogen).

Approximately 30,000 cells in suspension were placed in the coated culture dish. Upon resuspension, the culture medium was changed to CO_2_-independent DMEM (PN:12800-017; Invitrogen) with 4 mM NaHCO3 buffered with 20 *μ*M HEPES/NaOH (pH 7.2), for AFM experiments ∼2 hours before the measurement [17, 56–58].

The experimental setup included an AFM (Nanowizard I; JPK Instruments, Carpinteria, CA) that was mounted on a Zeiss Axiovert 200M optical, wide-field microscope using a 20x objective (Plan Apochromat, NA = 0.8; Zeiss, Oberkochen, Germany) along with a CCD camera (DMK 23U445 from The Imaging Source, Charlotte, NC). Cell culture dishes were kept in a petri-dish heater (JPK Instruments) at 37°C during the experiment. Before every experiment, the spring constant of the cantilever was calibrated by thermal noise analysis (built-in software; JPK) using a correction factor of 0.817 for rectangular cantilevers [59]. The cantilevers used were tipless, 200-350 *μ*m long, 35 *μ*m wide, and 2 *μ*m thick (CSC37, tipless, no aluminum; Mikromasch, Sofia, Bulgaria). The nominal force constants of the cantilevers ranged between 0.2 and 0.4 N/m. The cantilevers were supplied with a wedge, consisting of UV curing adhesive (Norland 63; Norland Products, East Windsor, NJ) to correct for the 10° tilt [60]. The measured force, piezo height, and time were output with a time resolution of at least 500 Hz.

#### Dynamic AFM-based cell confinement

Preceding every cell compression, the AFM cantilever was lowered to the dish bottom in the vicinity of the cell until it touched the surface and then retracted to ≈14 *μ*m above the surface. Subsequently, the free cantilever was moved and placed on top of the cell. Thereupon, a bright-field image of the equatorial plane of the confined cell was recorded to evaluate the equatorial radius *Req* at a defined cell height h. Cells were confined between dish bottom and cantilever wedge. Then, oscillatory height modulations of the AFM cantilever were carried out with oscillation amplitudes of 0.25 *μ*m at a frequency of 1 Hz.

During this procedure, the cell was on average kept at a normalized height *h/D* between 60 and 70%, where *D* = 2(3*/*(4*π*)*V*)^1*/*3^ and V is the estimated cell volume. Using molecular perturbation with cytoskeletal drugs, we could show in previous work that at these confinement levels, the resulting mechanical response of the cell measured in this setup is dominated by the actin cortex (see Figure 4 in [57] and Figure S7 in [5]). This is further corroborated by our observation from previous work that the smallest diameter of the ellipsoidal cell nucleus in suspended interphase cells is smaller than 60% of the cell diameter (see Figure S4 in [17]). Additionally, it has been shown that for a nucleus-based force response, when measuring cells in suspension with AFM, a confinement of more than 50% of the cell diameter is needed [61].

#### Data analysis

The data analysis procedure was described in detail in an earlier work [57]. In our analysis, the force response of the cell is translated into an effective cortical tension *γ* = *F/*[*A*_*con*_(1*/R*_1_ + 1*/R*_2_)], where *A*_*con*_ is the contact area between confined cell and AFM cantilever and *R*_1_ and *R*_2_ are the radii of principal curvatures of the free surface of the confined cell. Here *R*_1_ is estimated as half the cell height *h* and *R*_2_ is identified with the equatorial radius *R*_*eq*_ [17, 56, *57]*. *Cell height h* and equatorial radius *R*_*eq*_ were estimated from the AFM readout and optical imaging, respectively [34]. For the determination of the radius of the contact area *Acon*, see also Supplementary Section 5 in [62].

Oscillatory force and cantilever height readouts were analyzed in the following way: for every time point, effective cortical tension *γ* and surface area strain *ϵ*(*t*) = (*A*(*t*)− *< A >*)*/ < A >* were calculated. Here, *A*(*t*) is the total surface area of the confined cell. It is estimated as the area of a rotationally symmetric body with semi-circular free-standing side walls at cell height *h* and equatorial radius *Req*, i.e. 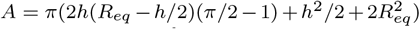 [17]. An amplitude and a phase angle associated to the oscillatory time variation of effective tension *γ* and surface area strain are extracted by sinusoidal fits. To estimate the value of the complex elastic modulus at a distinct frequency, we determine the phase angles *φγ* and *φϵ* as well as amplitudes *Aγ* and *Aϵ* of active cortical tension and surface area strain, respectively. The complex elastic modulus at this frequency is then calculated as *Aγ/Aϵ* exp(*i*(*φγ − φϵ*)).

Statistical analyses of cortex mechanical parameters were performed in MATLAB using the commands “boxplot” and “ranksum” to generate boxplots and determine p-values from a Mann-Whitney U-test (two tailed), respectively.

### 5.3. Plasmids and transfection

Transfection of cells was performed transiently with plasmid DNA using Turbofectin 8.0 (PN: TF81001, Origene), according to the manufacturer’s protocol. To achieve post-EMT conditions, MCF-7 cells were seeded at day -1. The cells were then transfected at day 0 and treated with 100 nM TPA (in the case of modMCF-7 cells). The Cells were then imaged at day 2. The plasmid MApple-LC-Myosin-N-7 was a gift from Michael Davidson (Addgene plasmid 54920; http://n2t.net/addgene : 54920; RRID : Addgene 54920). The plasmid MCherry-Actin-C-18 was a gift from Michael Davidson (Addgene plasmid 54967; http://n2t.net/addgene : 54967; RRID : Addgene 54967 [63]).

### 5.4. Imaging of transfected cells

Transfected cells were placed on PLL-g-PEG coated fluorodishes (FD35-100) with CO_2_-independent culture medium (described before). Cellular DNA was stained with Hoechst 33342 solution (PN:62249, Invitrogen) in order to distinguish between mitotic and interphase cells. During imaging, cells were maintained at 37°C using an Ibidi heating stage. Imaging was done using a Zeiss LSM700 confocal microscope of the CMCB light microscopy facility, incorporating a Zeiss C-Apochromat 40x/1.2 water objective. Images were taken at the equatorial diameter of each cell at the largest cross-sectional area (see Fig. 6a).

### 5.5. Calculation of cortex-to-cytoplasm ratios

This has been described before [5]. In short, using a MATLAB custom code, the cell boundary was identified (Fig. 1b shows an exemplary cell, the cell boundary is marked in red). Along this cell boundary, 200 radial, equidistant lines were determined by extending 1.5 *μ*m to the cell interior and 2.5 *μ*m into the exterior (Fig. 1b, red lines orthogonal to cell boundary, only every tenth line was plotted out of 200). The radial fluorescence profiles corresponding to these lines were averaged over all 200 lines (Fig. 1c, blue curve). This averaged intensity profile is then fitted by a linear combination of an error function (cytoplasmic contribution) and a skewed Gaussian curve (cortical contribution), see Fig. 1c, orange curve. The respective fit formula is given by

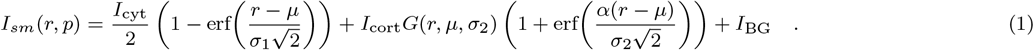

Here *p* = *{μ, σ*_1_, *σ*_2_, *I*_cyt_, *I*_cort_, *I*_BG_, *α}* are fit parameters, which determine the position of the cortex (*μ*), the slope of the error function decay (*σ*_1_), the width of the cortical Gaussian peak *σ*_2_, the amplitudes of the error function and the Gaussian peak (*I*_cyt_ and *I*_cort_) and the skewness of the Gaussian peak (*α*). To calculate the cortex-to-cytoplasm ratio, the fitted skewed Gaussian is integrated and the obtained integral is then normalized by the cytoplasmic intensity *Icyt* [5].

### 5.6. Immunostaining and confocal imaging of cells

The suspended (interphase or STC-arrested mitotic) cells were fixed with 3.7% PFA/PBS for 10 minutes (10% TCA for 15 minutes for Rac1, RhoA and RhoC, and -20°C ethanol for 10 minutes for Arp2) at room temperature, followed by a 10 min permeabilization step in 0.2% Triton X-100. The cells were then blocked for 1 hour at room temperature with 5%BSA/PBS. The cells were then treated with primary antibody of Rac1 (PN:PA1-091-X, Thermofisher), RhoA (PN:NBP2-22528, Novus Bio), RhoC (PN:GTX100546, GeneTex), CFL1 (PN:660571-1-lg, Proteintech), p-CFL1 (Ser3, PN:3313T, Cellsignalling), p-LIMK1 (Thr508, PN:E-AB-20918, Elabscience), mDia1 (PN:20624-1-AP, Proteintech) and Arp2 (PN:10922-1-AP, Proteintech) overnight at 4°C in 5%BSA/PBS. Cells were then treated with the corresponding secondary Alexa Fluor 488 conjugate at a concentration of 1:1000 in 5%BSA/PBS for 2 hours at room temperature. At the same time, cells were treated with 5 *μ*g/mL DAPI (2 minutes) and 0.2 *μ*g/mL Phalloidin-iFluor-647 (10 minutes) in 5% BSA/PBS solution. Images were taken with a Zeiss LSM700 confocal microscope of the CMCB light microscopy facility, incorporating a Zeiss C-Apochromat 40x/1.2 water objective. Images were taken at the equatorial diameter of each cell showing the largest cross-sectional area.

### 5.7. Western blotting

Protein expression in MCF-7 cells before and after EMT was analyzed using western blotting. Cells were seeded onto a 6-well plate and grown up to a confluency of 80-90% with or without EMT-inducing agents. Thereafter, cells were lysed in SDS sample/lysis buffer (62.5 mM TrisHcl pH 6.8, 2% SDS, 10% Glycerol, 50 mM DTT and 0.01%Bromophenolblue). For analysis of protein expression in mitotic cells, STC was added at a concentration of 2 *μ*M to the cell medium 12-18 hours before cells were harvested in order to enrich mitotic cells. For harvesting, mitotic cells were collected by shake-off and/or flushing of the medium.

Cell lysates were incubated for 30 minutes with the lysis buffer at 4°C. They were then boiled for 10 minutes. 10/20 *μ*L of the cleared lysate was then used for immunoblotting. The cleared lysates were first run on precast protein gels (PN:456-1096 or 456-1093, Bio-Rad) in MOPS SDS running buffer (B0001, Invitrogen). Subsequently, proteins were transferred to Nitrocellulose membranes (GE10600012, Sigma-Aldrich). Nitrocellulose membranes were blocked with 5% (w/v) skimmed milk powder (T145.1, Carl Roth, Karlsruhe, Germany) in TBST (20 mM/L Tris-HCl, 137 mM/L NaCl, 0.1% Tween 20 (pH 7.6)) or 5% (w/v) BSA for phospho-antibodies for 1 h at room temperature followed by washing with TBST, and incubation at 4°C overnight with the corresponding primary antibody diluted 1:500 (p-LIMK1), 1:1000 (mDia1, Arp2, CFL1 and p-CFL1) and 1:5000 (GAPDH) in 5% (w/v) bovine serum albumin/TBST solution. Thereupon, the blots were incubated with appropriate secondary antibodies conjugated to horseradish peroxidase, Goat anti-mouse HRP (PN: ab97023, Abcam) or Goat anti-rabbit HRP (PN: ab97051, Abcam) at 1:5000 dilution in 5% (w/v) skimmed milk powder in TBST for 1 h at room temperature. After TBST washings, specifically bound antibodies were detected using Pierce enhanced chemiluminescence substrate (ECL) (PN:32109, Invitrogen). The bands were visualized and analyzed using a CCD-based digital blot scanner, ImageQuant LAS4000 (GE Healthcare Europe, Freiburg, Germany). Primary antibodies used are as follows: GAPDH (PN:ab9485, Abcam), CFL1 (PN:660571-1-lg, Proteintech), p-CFL1 (PN:3313T, Cellsignalling), p-LIMK1 (PN:ELA-E-AB-20918, Biozol), mDia1 (PN:20624-1-AP, Proteintech) and Arp2 (PN:10922-1-AP, Proteintech). Two-tailed t-test (for two samples with unequal variance) was used for statistical analysis using Microsoft Excel.

### 5.8. Gene knock-down through RNA interference

Transfections were done targeting the genes CFL1 (siRNA, ID s2938, Thermofisher) or RHOC (esiRNA, HU-06597-1 Eupheria Biotech) at an RNA concentration of 25 nM, using the transfection reagent Lipofectamine RNAiMax (Invitrogen) according to the protocol of the manufacturer. Firefly luciferase esiRNA (FLUC, Eupheria Biotech) was used as a negative control, while EG5/KIF11 esiRNA (HU-01993-1, Eupheria Biotech) was used as a positive control. In all experiments, EG5/KIF11 caused mitotic arrest of more than 60-70% of the cells, showing a transfection efficiency of at least 60% in each experiment. Knock-down of RHOC and CFL1 was confirmed through Western blotting, see Fig. S2 and Fig. S3b-d.

For AFM measurements, at day -1, 30,000 cells were seeded into a 24-well plate (NuncMicroWell Plates with Nunclon; Thermo Fisher Scientific, Waltham, MA, USA). At day 0 the transfection was done. The transfected cells were imaged at day 2. For post-EMT conditions, the cells were kept in 100 nM TPA from day 0 to day 2. For mitotic cells, ≈12-24 hours before measurements, cells were detached, diluted, and transferred onto a glass-bottom Petri dish (FD35-100, World Precision Instruments) with 2 *μ*M STC added ≈2 hours before measurement. For interphase cells, 1-2 hours before measurement the cells were detached and transferred to PLL-g-PEG-coated Petri dishes (see Section on AFM Measurements of Cells).

For Western blotting, at day -1, 800,000 cells were seeded into a 6-well plate (NuncMicroWell Plates with Nunclon; ThermoFisher Scientific, Waltham, MA, USA). At day 0 the transfection was done. The transfected cells were then lysed at day 2 as described in the Western blotting section. For post-EMT conditions, the cells were kept in 100 nM TPA from day 0 to day 2. For mitotic cells, ≈12-18 hours before lysing, 2 *μ*M STC was added.

## Author Contributions

K. H. and A. F. performed the experiments. K. H. and E. F.-F. designed the experiments. K. H. performed data analysis. K. H. and E. F.-F. wrote the manuscript.

## Conflict of interest

There are no conflicts to declare.

## Acknowledgments

EFF acknowledges financial support from the Deutsche Forschungsgemeinschaft under Germany’s Excellence Strategy, EXC-2068-390729961, Cluster of Excellence Physics of Life of TU Dresden. Furthermore, EFF was funded by the Deutsche Forschungsgemeinschaft (DFG, German Research Foundation) – project number 495224622 (FI 2260/8-1) and by the grant FI 2260/7-1. In addition, the authors thank the CMCB and PoL Light Microscopy Facility for excellent support.

## Notes

### Competing Interest Statement

The authors have declared no competing interest.

### Summary of Updates

manuscript text

