## Supplementary text for "EMT induces cell-cycle-dependent changes of Rho GTPases and downstream effectors"

Kamran Hosseini,[1,2] Annika Frenzel,[1,2] and Elisabeth Fischer-Friedrich[1,2,3,a)]
[1)] *Cluster of Excellence Physics of Life, Technische Universität Dresden, Dresden, Germany*
[2)] *Biotechnology Center, Technische Universität Dresden, Dresden, Germany*
[3)] *Faculty of Physics, Technische Universität Dresden, Dresden, Germany*

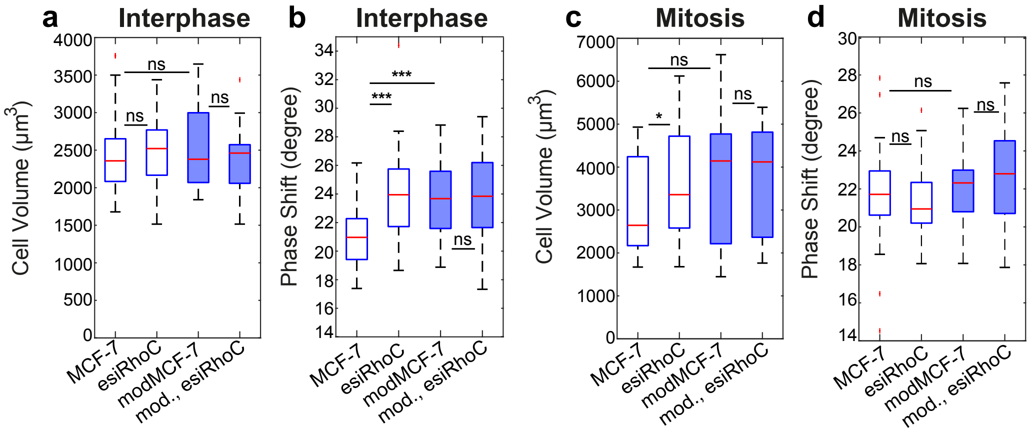


Figure S1. Changes of cell volume and of the phase shift of the complex elastic modulus of the cortex upon RhoC-knockdown in MCF-7 cells in pre- and post-EMT conditions in suspended interphase cells (a-b) and cells in mitotic arrest (c-d). Data correspond to measurement results presented in Fig. 1i-l, main text. Post-EMT cells are referred to as modMCF-7. Number of cells measured: a-b: MCF-7 n=39, esiRhoC n=37, modMCF-7 n=37, esiRhoC n=39, c-d: MCF-7 n=27, esiRhoC n=29, modMCF-7 n=29, esiRhoC n=28. n.s.: $p > 0.05, * : p < 0.05$, $** : p < 0.01$, $*** : p < 0.001$.

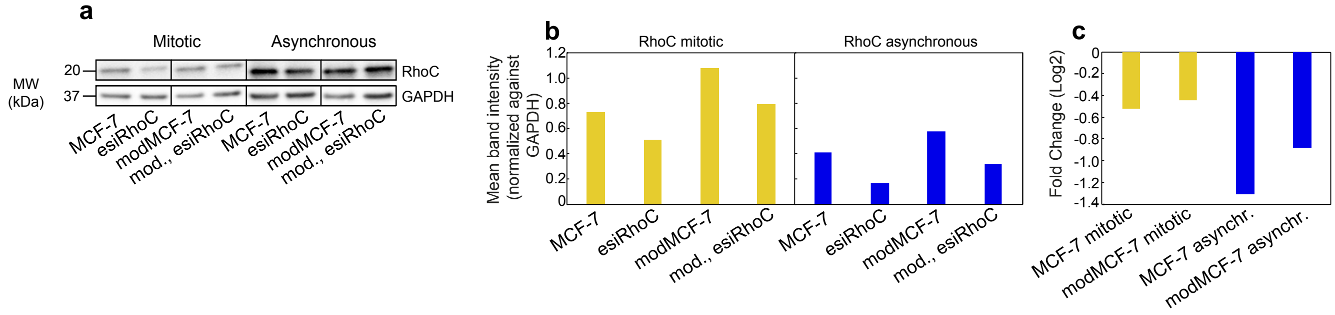


Figure S2. Relative changes of RhoC abundance upon knock-down. a-c) Western blots and quantification showing RhoC abundance in lysates of MCF-7 and modMCF-7 (in mitotic and asynchronous cell populations) in control conditions and upon RhoC knock-down. Knock-down was achieved through RNA interference. Normalization was done against GAPDH. The Western blots show successful expression changes of the target protein upon knock-down.

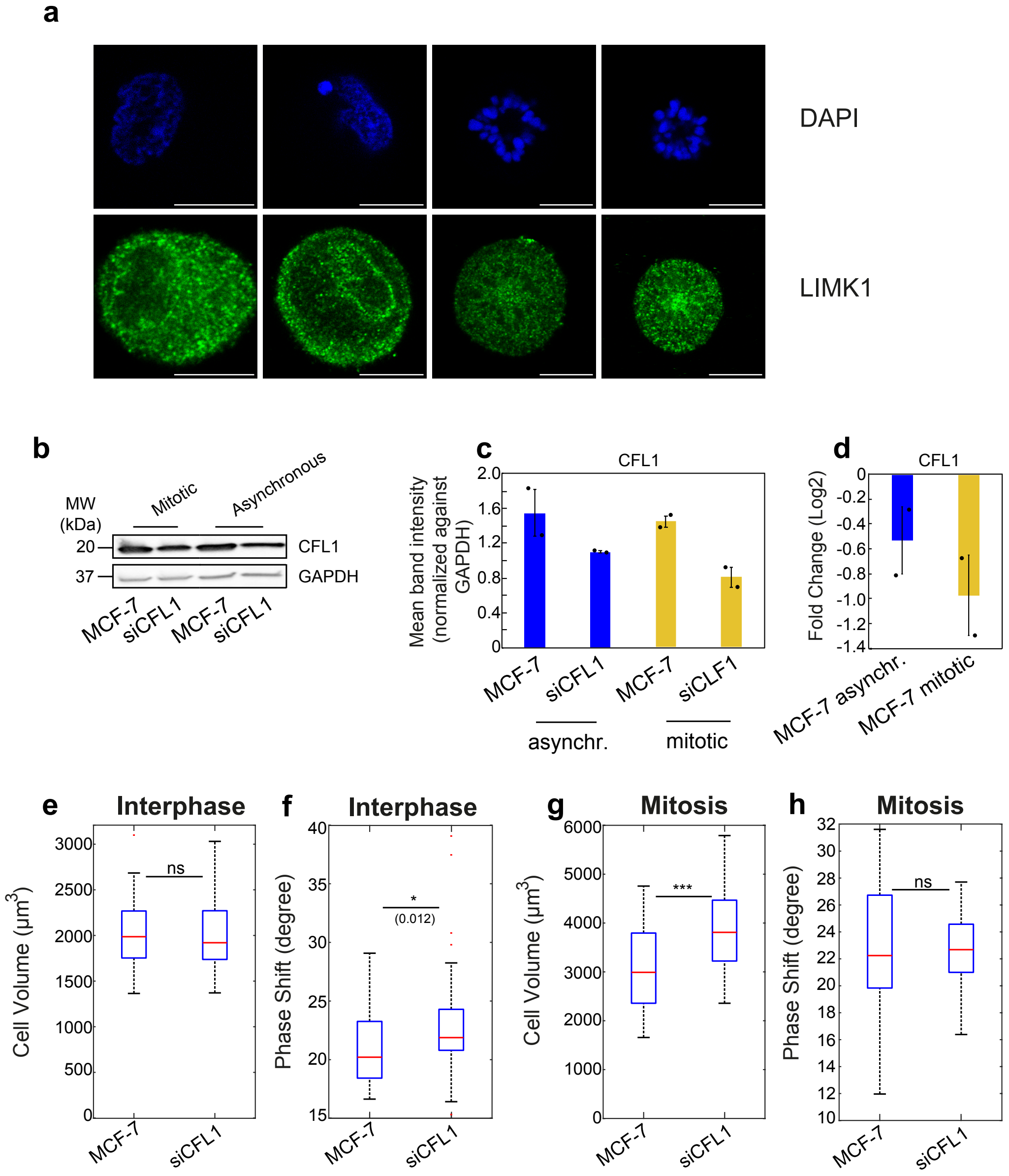


Figure S3. a) Representative confocal images of Limk1 (immunostained, green) and DNA (DAPI, blue) in suspended interphase cells and STC-arrested mitotic cells in pre-EMT (MCF-7) and post-EMT (modMCF-7) conditions. Scale bar: 10 $\mu$m. b-d) Western blots and quantification showing CFL1 abundance in lysates of MCF-7 (in mitotic and asynchronous cell populations) in control conditions and upon CFL1 knock-down. Knock-down was achieved through RNA interference. Normalization was done against GAPDH. Individual data points are depicted in black. Error bars represent standard error of the mean. The Western blots show successful expression changes of the target protein upon knock-down. e-h) Cell volume and phase shift changes upon CFL1-knockdown MCF-7 cells corresponding to measurements presented in Fig. 3f-i, main text, for suspended interphase cells (e-f) and cells in mitotic arrest (g-h). Number of cells measured: e-f: MCF-7 n=49, siCFL1 n=49, g-h: MCF-7 n=46, siCFL1 n=50. n.s.: $p > 0.05, * : p < 0.05$, $** : p < 0.01$, $*** : p < 0.001$.

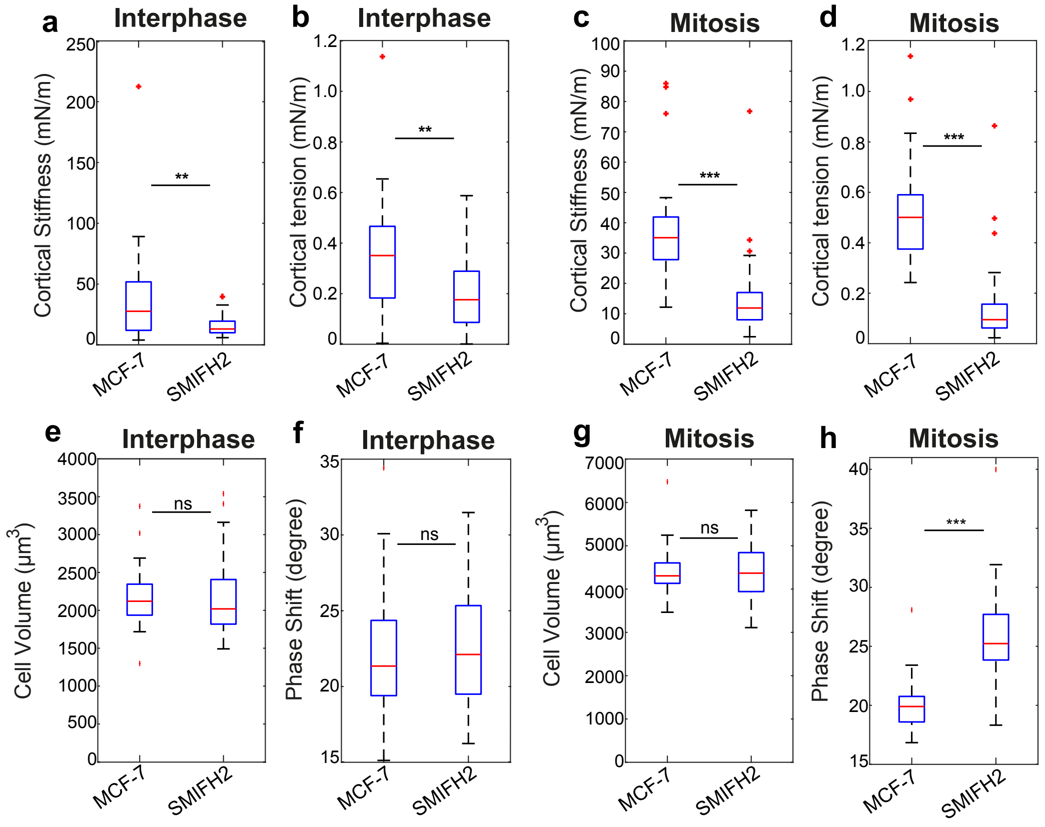


Figure S4. a-d) Formin inhibition using 40 $\mu$M SMIFH2 elicits cortical softening and tension reduction in the actin cortex in interphase (a-b) and mitotic MCF-7 cells (c-d). e-h) Corresponding cell volume and phase shift changes upon formin inhibition for suspended interphase cells (e-f) and cells in mitotic arrest (g-h). Number of cells measured: a-b, e-f: MCF-7 n=26, SMIFH2 n=21, c-d, g-h: MCF-7 n=32, SMIFH2 n=32. Measurements represent at least two independent experiments. n.s.: $p > 0.05, * : p < 0.05$, $** : p < 0.01$, $*** : p < 0.001$.

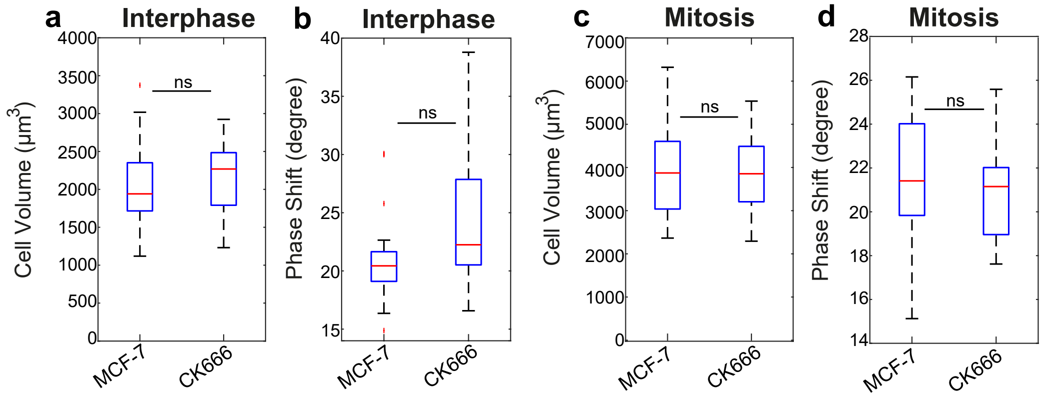


Figure S5. Cell volume and phase shift changes upon Arp2/3 inhibition in MCF-7 cells, using 50 $\mu$M CK666, corresponding to measurements presented in Fig. 5f-i, main text, for suspended interphase cells (a-b) and cells in mitotic arrest (c-d). Number of cells measured: a-b: MCF-7 n=24, CK666 n=24, c-d: MCF-7 n=27, CK666 n=24. n.s.: $p > 0.05, * : p < 0.05$, $** : p < 0.01$, $*** : p < 0.001$.

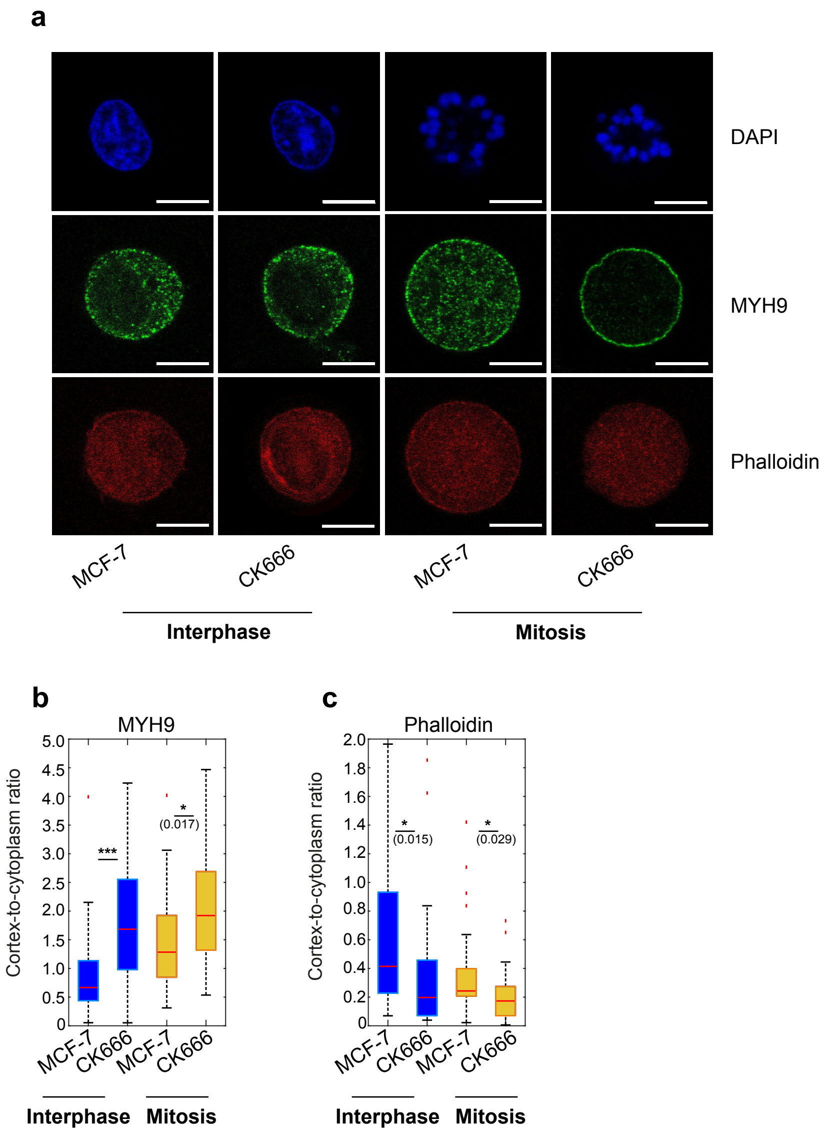


Figure S6. Arp2/3 activity affects myosin localization to the cortex. a) Representative confocal images of suspended interphase cells and STC-arrested mitotic MCF-7 cells in control conditions and with Arp2/3 inhibition via CK666. Cells were fixed and DAPI-stained for DNA (blue), Phalloidin-stained for actin (red) and immunostained for MYH9 (green), see Materials and Methods. Scale bar: 10 $\mu$m. b-c) Cortex-to-cytoplasm ratio of MYH9 (b) and actin (c) inferred from images as shown in panel a before and after Arp2/3 inhibition with 50 $\mu$M CK666. Blue-shaded boxplots represent suspended interphase cells and yellow-shaded boxplots represent STC-arrested mitotic cells. Number of cells analyzed: b: interphase: MCF-7 n=43, CK666 n=43, mitosis: MCF-7 n=29, CK666 n=30, c: interphase: MCF-7 n=22, CK666 n=23, mitosis: MCF-7 n=30, CK666 n=30. n.s.: $p > 0.05, * : p < 0.05$, $** : p < 0.01$, $*** : p < 0.001$.
